# Novel polygenic risk score as a translational tool linking depression-related changes in the corticolimbic transcriptome with neural face processing and anhedonic symptoms

**DOI:** 10.1101/556852

**Authors:** Klara Mareckova, Colin Hawco, Fernanda C. Dos Santos, Arin Bakht, Navona Calarco, Amy E. Miles, Aristotle N. Voineskos, Etienne Sibille, Ahmad R. Hariri, Yuliya.S. Nikolova

## Abstract

Convergent data from imaging and postmortem brain transcriptome studies implicate corticolimbic circuit (CLC) dysregulation in the pathophysiology of depression. To more directly bridge these lines of work, we generated a novel transcriptome-based polygenic risk score (T-PRS), capturing subtle shifts towards depression-like gene expression patterns in key CLC regions, and mapped this T-PRS onto brain function and related depressive symptoms in a non-clinical sample of 478 young adults (225 men; age 19.79+/−1.24) from the Duke Neurogenetics Study. First, T-PRS was generated based on common functional SNPs shifting CLC gene expression towards a depression-like state. Next, we used multivariate partial least squares regression to map T-PRS onto whole-brain activity patterns during perceptual processing of social stimuli (i.e., human faces). For validation, we conducted a comparative analysis with a PRS summarizing depression risk variants identified by the Psychiatric Genomics Consortium (PGC-PRS). Sex was modeled as moderating factor. We showed that T-PRS was associated with widespread reductions in neural response to neutral faces in women and to emotional faces and shapes in men (multivariate p<0.01). This female-specific reductions in neural response to neutral faces was also associated with PGC-PRS (multivariate p<0.03). Reduced reactivity to neutral faces was further associated with increased self-reported anhedonia. We conclude that women with functional alleles mimicking the postmortem transcriptomic CLC signature of depression have blunted neural activity to social stimuli, which may be expressed as higher anhedonia.

## INTRODUCTION

Major depressive disorder (MDD or depression) is the leading cause of disability worldwide^1^, with lifetime prevalence up to 17%^2^. Studies on the biological basis of depression using in vivo neuroimaging or postmortem gene expression often focus on the same neural circuits, but have largely run in parallel, precluding an integrative understanding of this complex and heterogeneous disease. In vivo neuroimaging^3–10^ and postmortem transcriptome studies^11^ have nonetheless independently converged to associate depression with dysregulation in a distributed corticolimbic circuit (CLC), which lies at the intersection of affective, cognitive, and perceptual processing.

Functional neuroimaging studies of the CLC in depression have typically focused on the amygdala as a primary region of interest, because this structure serves as a processing hub through which associative learning of threat and related physiological and behavioral responses are coordinated^3, 5, 6^. These studies associate depression with increased amygdala reactivity to threat^3^, which may pre-date the development of depressive symptoms^4^ and persist after remission^3, 5, 6^. Consistent with the heterogeneity of the disease, however, other studies have reported blunted amygdala reactivity in individuals with, or at risk for, depression^7–10, 12^. Although the behavioral implications of these divergent neural phenotypes are incompletely understood, amygdala hyper-reactivity has generally been interpreted as reflecting a depression risk pathway associated with high anxiety and stress sensitivity^4^, while hypo-activity may reflect reduced behavioral engagement, consistent with co-occurring trait-like psychomotor slowing^7, 10, 12^ and motivational deficits^10^. Functional connectivity studies using both hypothesis- and data-driven approaches have additionally reported reduced coupling between amygdala and prefrontal regulatory regions^13, 14^, as well as broader dysconnectivity within and beyond the CLC^15–20^.

In parallel to this functional neuroimaging work, postmortem studies in depression have demonstrated changes in gene expression patterns in key CLC nodes, which may partially explain the observed macroscale CLC dysregulation noted above. A recent meta-analysis of 8 transcriptome datasets^11^ identified 566 genes consistently up- or down-regulated in individuals with depression across the 3 amygdala, subgenual anterior cingulate (sgACC), and dorsolateral prefrontal cortex (dlPFC)^11^. The altered molecular pathways suggested by these transcriptomic findings include reduced neurotrophic support, neuronal signaling, and GABA function. These pathways have also been implicated in normal lifelong aging of the brain^21, 22^, lending support to the notion that depression may be associated with accelerated, or anticipated, molecular brain aging^21, 23^. Finally, functional neuroimaging studies suggest normal aging is associated with blunted CLC processing of emotional faces^24^, mirroring neural activity patterns observed in specific depression subtypes^7–10, 12^.

Despite this promising convergence, postmortem findings have limited clinical utility, as they provide little insight into the link between gene expression changes in the CLC and in vivo CLC reactivity or any clinically relevant pre-mortem symptoms. To address this important translational gap, we developed a novel transcriptome-based polygenic risk score (T-PRS) comprised of common gene expression-altering variants, partially mimicking the postmortem depression CLC transcritpome. Using a data-driven multivariate approach (partia least squares - PLS regression), we then mapped this score onto fMRI-assessed neural processing of social stimuli in a large sample of young adults participating in the Duke Neurogenetics Study.

In a subsequent validation analysis, we compared the effects of this novel T-PRS to those of a non-overlapping PRS based on variants identified using more established polygenic risk scoring techniques derived from case-control genome-wide association studies by the Psychiatric Genomics Consortium MDD working group (PGC-PRS^25^). Given well-documented sex differences in depression risk^26–30^, and CLC function^31–34^, sex was included as a moderating factor in all analyses. We hypothesized that both PRS will be associated with heightened neural responses to threatening faces and increased depressive symptoms.

## MATERIALS AND METHODS

### Participants

The study used archival data from 482 young adult university students (226 men, 256 women; mean age 19.78 +− 1.23) who successfully completed the Duke Neurogenetics Study (DNS). All participants provided informed consent in accordance with Duke University guidelines prior to participation and were in good general health (see Nikolova et al^35^ for full exclusionary criteria). Participants were screened for DSM-IV Axis I and select Axis II disorders (Antisocial and Borderline Personality Disorder) using the eMINI^36^, but a current or lifetime diagnosis of a disorder was not exclusionary (**Supplementary Table 1**). The Duke University Institutional Review Board approved all study procedures.

We limited our analyses to Non-Hispanic Caucasians to match the ethnic background of the postmortem cohorts used to develop our T-PRS and corrected for the resulting population structure as follows: The presence of relatedness in the sample was assessed through proportional identity by descent (PIHAT). In pairs of individuals showing a PIHAT higher than 0.2, the individual showing higher genotype missingness (lower CR) was excluded from analysis. In order to avoid spurious associations due to population stratification, after filtering by self-reported ethnicity we further assessed any residual stratification using a Multidimensional scaling (MDS) and clustering method implemented in PLINK v1.9^37^, following the thresholds and recommendations published elsewhere^38, 39^. Briefly, genetic data of 629 individuals from known different ethnic backgrounds (a reference panel from 1000 Genomes Project) was assessed and used as reference panel for ethnicity. Two individuals from current sample that did not cluster with the 1000 genomes Europeans were excluded from analysis (**Supplementary Figure 1**). MDS analysis was performed again in the final dataset (n=478; 225 men, 253 women; mean age 19.79 +− 1.24), used also for all further analyses, to generate 10 principal components. The first component (C1) alone was able to explain more than 40% of the variance, therefore it was included in all analyses as a covariate correcting for residual population substructure^38^ (**Supplementary Figure 2**).

### Calculation of the transcriptome-based polygenic risk score

DNA extraction and genotyping was performed as previously described in Nikolova et al^10^. A transcriptome-based polygenic risk score (T-PRS) characterizing depression-related gene expression changes in the CLC was developed based on a list of 566 genes generated by a prior meta-analysis of case-control postmortem brain transcriptome datasets (n=101 postmortem subjects; 50 MDD, 51 controls^11^). Relying on the GTEx “cortex” tissue as a reference transcriptome, the PrediXcan tool^40^ successfully “imputed” relative cortical expression levels on the individual participant level for 76 out of these 566 genes based on peripheral SNPs assessed in the DNS (see Figure 1). Notably, expression levels for the remaining genes were not imputed because these genes did not have significant cis-eQTLs identified in the reference tissue, possibly due to lower expression heritability^40^. A PRS was created as a sum of the 76 imputed expression values (normalized to a consistent scale across genes) with higher T-PRS reflecting a more depression-like CLC transcriptome. To allow easy computation of our novel T-PRS scores in other studies, we provide a list of the 76 genes in **Supplementary Table 2**.

**Figure 1.**
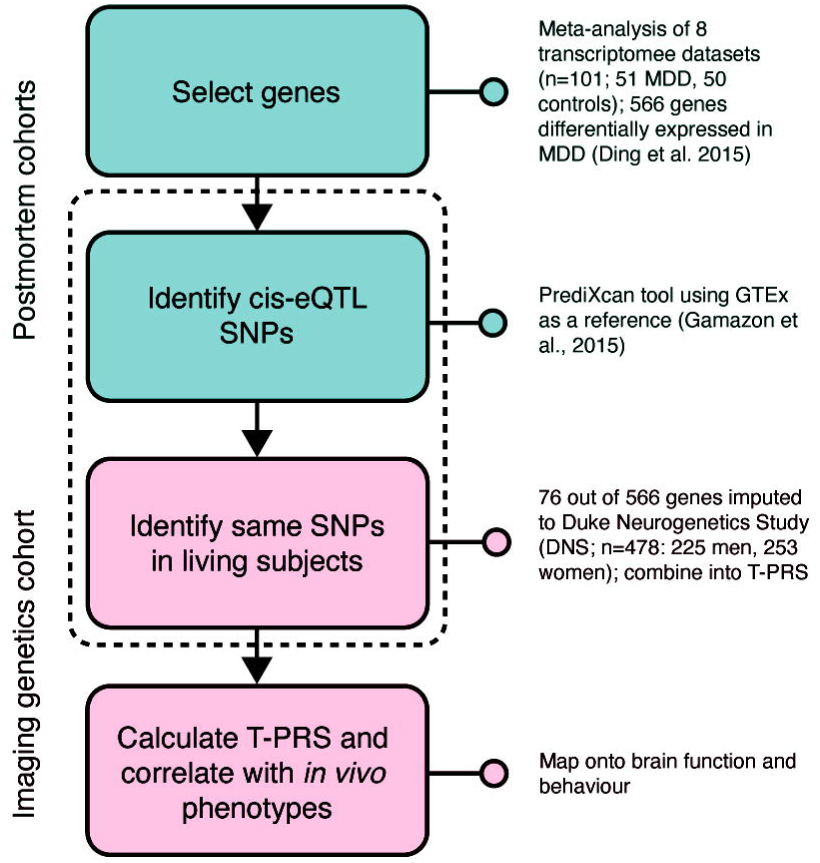
Study design. A list of 566 genes associated with depression on the transcriptome-wide level was selected based on a prior meta-analysis of 8 postmortem brain transcriptome datasets (Ding et al., 2015). Building on reference data from the combined genomic-transcriptomic GTEx database, the PrediXcan tool, was then used to “impute” cortical expression of 76 of these genes based on 478 young adults participating in the Duke Neurogenetics Study (DNS). The imputed values for each of these genes were weighted by their original association with depression and summed into a polygenic risk score (T-PRS), such that higher values indicated a more depression-like transcriptome. This novel T-PRS was then mapped onto brain function during face processing.

### Calculation of the PGC-based polygenic risk score

Summary statistics from Wray et al.^25^ including genome-wide SNPs from the meta-analysis excluding 23andMe was downloaded from the PGC website and QC cleaned. Cleaning included checking the effect allele, confirming the genome build (NCBI Build 37/UCSC hg19, same as DNS sample); filtering effect variants by minimum allelic frequency (MAF>0.01) and quality of imputation (INFO>0.80). No ambiguous SNPs were identified in the current sample [rs34215985, rs115507122 and rs62099069 previously identified as ambiguous were not genotyped or imputed in the current sample]. There were no duplicated SNPs in this dataset and no sex chromosome variants reported. There is also no sample overlap or related individuals between PGC and DNS sample. PGC-PRS was calculated using the PLINK v1.9^38^ --score function and multiple p-value thresholds (5e10-8, 1e10e-6, 1e10-4, 1e10-3, 1e10-2, 0.01, 0.05, 0.5 and 1). DNS data was clumped according to the p-values in the Summary statistics following the parameters of significance threshold for index SNPs of 1, an LD threshold of 0.1 and a physical distance of 250kb between the index and the SNPs to be clumped. Using this strategy, 3,979,908 variants out of 6,372,415 variants were missing in the main dataset (13 loci from the 44 significant in Wray 2018 paper), and 166,296 clumps were formed and used to calculate the PRS. Total number of variants included in each score is indicated in **Supplementary Table 3**.

### Self-report measures

The short form of the Mood and Anxiety Symptom Questionnaire (MASQ)^41^ was used to provide information on general distress symptoms shared between depression and anxiety (general distress depression and anxiety scales; GDD and GDA, respectively) or symptoms unique to anxiety (anxious arousal, AA subscale) or depression (anhedonia, AD subscale)^41^. Early and recent life stress were assessed using the Childhood Trauma Questionnaire (CTQ^42^) and the Life Events Scale for Students (LESS^43^), respectively.

### Acquisition of fMRI data

BOLD fMRI data were acquired during a face-matching task^44^ at the Duke-UNC Brain Imaging and Analysis Center using one of two identical research-dedicated General Electric MR750 3T scanners. Briefly, participants were presented with blocks of neutral and emotional faces, interleaved with blocks of simple geometric shapes as a control condition, and were asked to indicate which one of two faces/shapes shown at the bottom of the screen was identical with a target face/shape shown at the top. Detailed information regarding the face-matching task as well as the acquisition protocol are provided in the Supplementary Methods.

### Multivariate whole-brain analysis

Partial Least Square (PLS) regression^45, 46^ was used to identify the functional impact of the T-PRS at the whole-brain level. PLS regression combines features of principal component analysis and multiple regression and is a multivariate approach that allows the simultaneous mapping of a multidimensional set of independent predictors (i.e. the relationship with polygenic risk and brain activity for each condition, separately for males and females) onto a multidimensional set of intercorrelated outcome variables (i.e., the spatial pattern of voxel loadings voxel time series across the whole brain). This is 7 achieved through the identification of a smaller set of latent variables (LVs), which best capture covariance between the independent variables and distributed spatial patterns of neural activity, without the need for a priori selected contrasts across experimental conditions or regions of interest^45^. The PLS thus identified LVs representing a spatial pattern of brain regions which share a common relationship between polygenic risk and brain activity in each condition (emotional faces, neutral faces, and shapes), separately by group (males/females). The LVs are similar to the components derived via principal component analysis (though optimized to detect relationships across a set of predictor variables); the number of LVs is determined by the input data and LVs explain progressively less covariance between the predictors.

In order to assess the statistical relevance of the LVs, a permutation approach was employed, rerunning the PLS while randomly resampling the order of conditions across 500 permutations to create a null distributions of saliences across LVs. This approach can be used to assess significance. As the permutation test works on the whole PLS model, it is not then necessary to correct for multiple comparisons for individual LVs.

Following LV identification, a bootstrapping procedure with 1000 iterations was used to test which specific voxels are reliably related to each significant LV. A bootstrap ratio for each voxel was calculated as the voxel salience divided by its bootstrap standard error. A bootstrap ratio of 2.5 (corresponding to >95% reliability) was used to threshold for all voxel pattern maps in PLS, as previously described^47^. A parallel analysis using the same parameters was conducted for all PGC-PRS to examine convergence between our novel T-PRS and a depression PRS generated based on more conventional polygenic risk modeling methods. PLS analyses involving the nine distinct PGC-PRS were corrected for multiple comparisons using a Bonferroni correction, wherein the significance threshold for an LV identified in each individual PGC-PRS analysis was set to p_corr_=0.05/9=0.005.

Finally, we conducted additional exploratory analyses linking PRS, multivariate brain activity patterns and symptoms. To this end, we extracted individual-level “brainscores” for the significant conditions, 8 reflecting subject-specific fit to multivariate brain activation patterns, and correlated them with each of the MASQ scales. This analysis was undertaken to examine whether the degree to which an individual participant fit the PRS-associated brain activity pattern was associated with symptoms measured independently (i.e., the clusters are derived independently of symptoms). We opted for this approach, rather than including PRS and symptoms in the same multivariate model, as we were primarily interested in validating the potential behavioral and clinical relevance of pre-identified genetically driven alterations in brain response, rather than identifying novel brain patterns where genetic and symptom effects converge. Further details regarding the PLS regression analysis are provided in the **Supplementary Methods**. Posthoc univariate analyses focused on the amygdala were conducted for continuity with prior work (see **Supplementary Methods**).

## RESULTS

### Effects of T-PRS on whole-brain activation

Our multivariate PLS analysis identified one significant latent variable (LV1; p<0.01), which accounted for 39.27% of crossblock covariance (i.e., shared variance between T-PRS, task condition, and voxel-wise brain activity). The spatial pattern of brain activity in this LV mapped onto 29 clusters greater than 20 voxels, spanning a distributed network including the mid-frontal and frontal superior regions, supplementary motor area, hippocampus, caudate, thalamus and cerebellum (see **Supplementary Table 4**). Contrary to our hypothesis, activity in all clusters was negatively correlated with the Neutral faces condition in women, indicating that, across this network, higher T-PRS was associated with female-specific blunting of activity driven by the neutral faces condition. In men, higher T-PRS was related to higher response in these clusters during the Emotional faces and Shapes conditions (Figure 2A).

**Figure 2.**
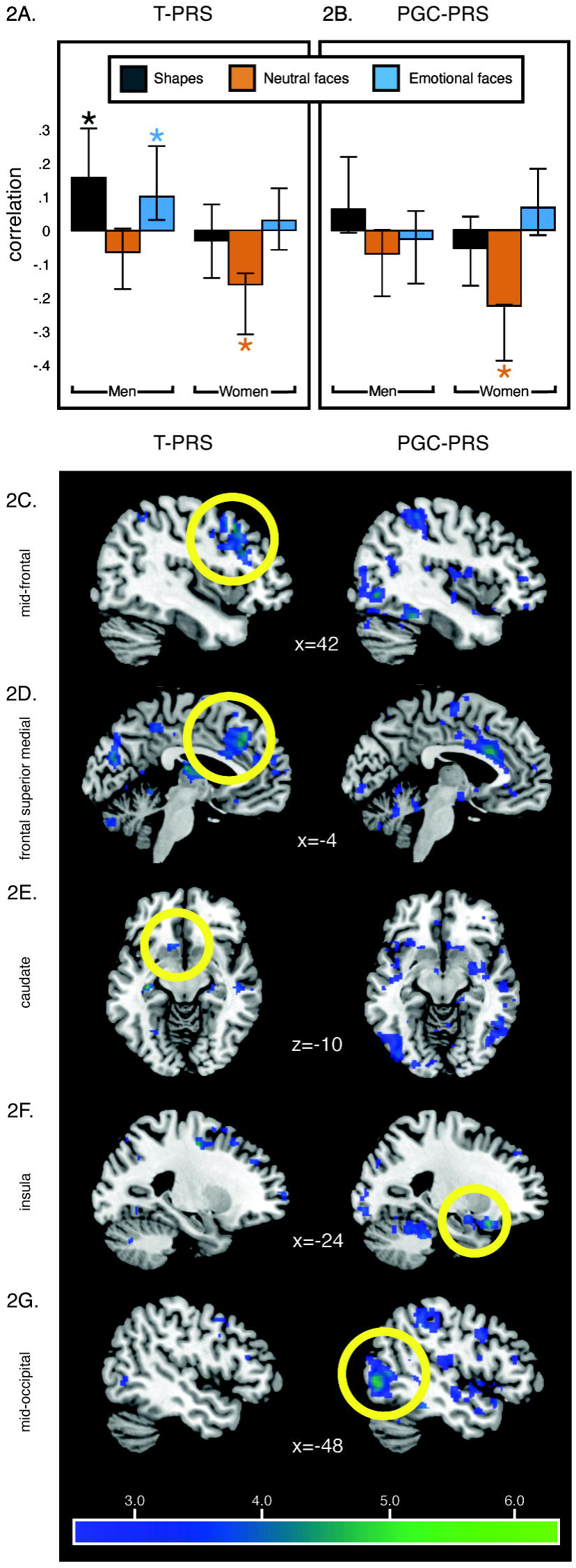
Brain response to social stimuli associated with the novel T-PRS and the PGC-PRS. Higher levels of T-PRS were associated with lower activity in the LV1 clusters during the Neutral faces condition in women and higher activity in the LV1 clusters during the Emotional faces and Shapes conditions in men (**2A**). Higher levels of PGC-PRS were associated with lower activity in the LV1 clusters during the Neutral faces condition in women (**2B**). Error bars represent 95% confidence intervals based on the bootstrapping distribution, which can be asymmetrical. Specifically, higher levels of T-PRS were more strongly associated with lower activity in frontal regions, including the mid-frontal (**2C**) and frontal superior medial (**2D**), and caudate (**2E**) and higher levels of PGC-PRS were associated with low activity in insula (**2F**) and mid-occipital (**2G**). These clusters survived the 2.5 bootstrap ratio (corresponding to 95% reliability) and were greater than 20 voxels.

### Validation and comparison to PGC-PRS

For comparison, we used the same data-driven multivariate framework to examine the whole-brain activity patterns associated with a series of PGC-PRS, all of which were uncorrelated with T-PRS (r values <03, p values > 0.100). Across thresholds, PLS analyses involving PGC-PRS identified a single 9 significant LV (**Supplementary Table 3**). Only the LVs identified by the analyses involving PGC-PRS computed at the p_gwas_<0.001 and p_gwas_<0.01 thresholds resulted in an LV whose significance survived correction at the adjusted p_corr_<0.005 threshold. Out of those, the score computed at p_GWAS_ <0.001 fit the imaging data better, as reflected in the higher cross-block covariance associated with its respective model, and was thus selected for visualization and interpretation. Specifically, this analysis identified one significant LV (p=0.04) which accounted for 37.92% of crossblock covariance and mapped onto 94 clusters greater than 20 voxels spanning anterior cingulum, amygdala, insula, postcentral, temporal, occipital regions and cerebellum (see **Supplementary Table 5**). Similarly to our T-PRS results, the best-fit PGC-PRS was associated with blunted reactivity to neutral faces in women (Figure 2B). Given the lack of direct correlation between the two PRS, this promising convergence of downstream effects provides additional confidence in the T-PRS effects, despite the unexpected condition and direction they emerged in.

Notably, both scores were associated with blunted activity to faces in key CLC nodes such as the hippocampus, as well as other regions spanning the mid-temporal, supplementary motor area and cerebelum. However, while the T-PRS was more strongly and specifically associated with reduced activity in the frontal regions and caudate (Figure 2C-E), the PGC-PRS effects were more strongly associated with reduced activity in insula and mid-occipital (Figure 2F-G). The results did not change substantially when the analysis was re-run with PRS score residualized for C1 (**Supplementary Figure 3**).

### Association with mood and anxiety symptoms

Additional exploratory analyses correlated PLS-derived “brainscores” for the significant conditions (brain response to neutral faces associated with T-PRS and PGC-PRS in women and brain response to emotional faces and shapes associated with T-PRS in men; viz Figure 2A), reflecting the degree of fit to genetic-associated brain activity patterns, with each of the four MASQ subscales. Brainscores reflecting T-PRS-associated blunting of neural response to neutral faces were associated with higher anhedonia in women (R^2^=0.03, p=0.007; Figure 3A), but had no effect on other MASQ facets.

**Figure 3.**
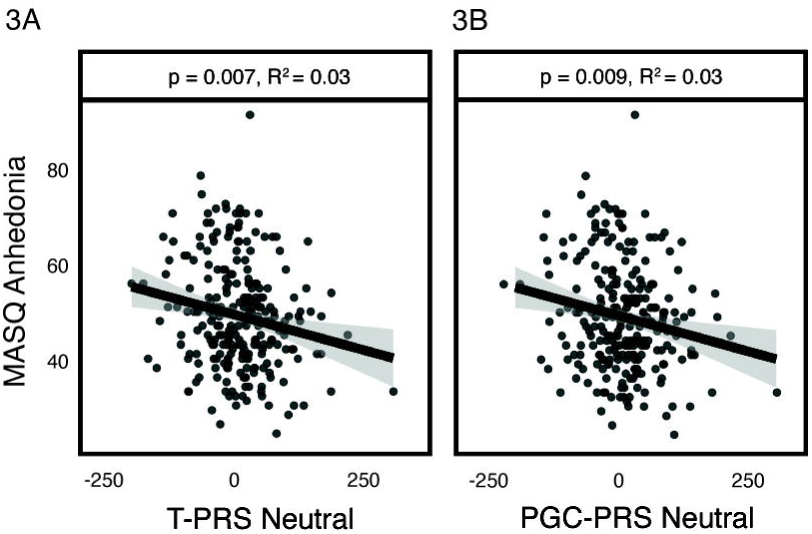
Brainscores and anhedonia. Greater blunting of the T-PRS LV1 regions was associated with more anhedonia in females (R^2^=0.03, p=0.007; **3A**).Similar negative relationship was also observed in females between the PGC-PRS-associated blunting of neural response to neutral faces and anhedonia (R^2^=0.03, p=0.009; Figure 3B).

Moreover, this association between T-PRS-associated blunting of neural response to neutral faces and anhedonia in women was independent of the remaining three MASQ domains, which capture symptoms unique to anxiety (anxious arousal) or shared between depression and anxiety (general distress scales), as well as early or recent life stress (beta=-0.14, p=0.006). Similar negative relationship was also observed in females between the PGC-PRS-associated blunting of neural response to neutral faces and anhedonia (R^2^=0.03, p=0.009; Figure 3B). Again, this association was independent of the remaining three MASQ domains, which capture symptoms unique to anxiety (anxious arousal) or shared between depression and anxiety (general distress scales), as well as early or recent life stress (beta=0.12, p=0.013). No additional associations with self-report symptoms emerged for the emotional faces or the shapes condition in men (all p>0.64).

Finally, a moderated mediation analysis revealed that brain response to neutral faces in the T-PRS-based LV1 regions mediated the relationship between T-PRS and anhedonia in women (ab=0.093, SE=0.047, 95% CI [0.019; 0.198]; Figure 4) but not men (ab=0.008, SE=0.033, 95% CI [-0.061; 0.076]). In contrast, the brain response to neutral faces in the PGC-PRS-based LV1 regions did not mediate the relationship between PGC-PRS and anhedonia in neither women (ab=-0.007, SE=0.005, 95% CI [-0.019; 0.002]), nor men (ab=-0.001, SE=0.003, 95% [-0.008; 0.004]).

**Figure 4.**
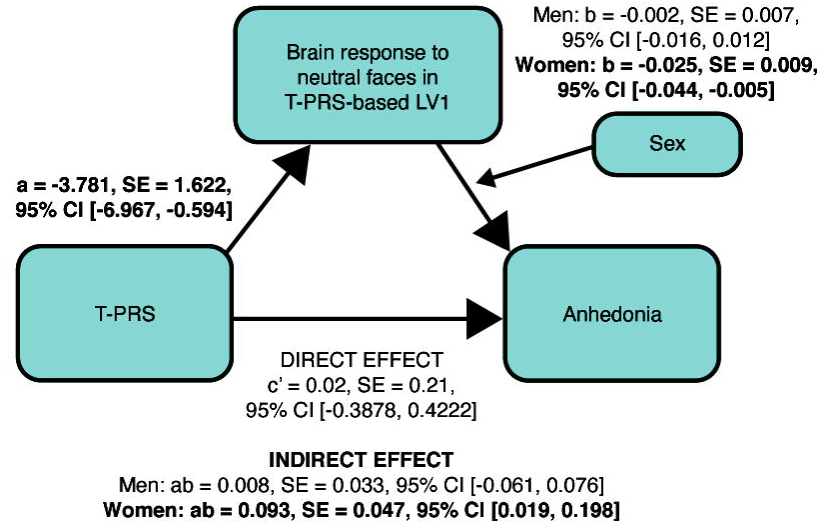
Brain response to neutral faces in T-PRS-based LV1 regions mediated the relationship between T-PRS and anhedonia in females. (ab=0.093, SE=0.047, 95% CI [0.019; 0.198]) **but not males** (ab=0.008, SE=0.033, 95% CI [-0.061; 0.076]).

For consistency with prior work, we also conducted a GLM focusing on amygdala reactivity to threatening and neutral faces vs shapes. These analyses confirmed the relationship bewteen higher T-PRS and more blunted neural response during observation of neutral faces (**Supplementary Figure 4**) and this relationship remained significant also after the correction for all covariates (sex, age, C1, early and recent stress, MDD diagnosis). Moreover, amygdala reactivity during observation of neutral faces (vs. shapes) showed a relationship with anhedonia (**Supplementary Figure 5**) and mediated the relationship between T-PRS and anhedonia (see **Supplementary Results** and **Supplementary Figure 6**).

## DISCUSSION

Here, we provide novel evidence that a polygenic score comprised of functional variants mimicking the postmortem transcriptomic signature of depression in key corticolimbic brain regions (T-PRS) is associated with female-specific widespread reductions in activity across cortex, subcortex, and cerebellum to neutral faces, and indirectly, with increased anhedonia, in a non-clinical sample of young adults. To validate these results, we demonstrated that similar female-specific blunting of neural reactivity to neutral faces was associated with an independent polygenic score derived using more traditional polygenic risk modeling approaches (i.e. non-overlapping genome-wide supported risk variants for depression; PGC-PRS).

This proof-of-principle study demonstrates that transcriptome-based polygenic risk scores may serve as a translational bridge between postmortem molecular data and in vivo neuroimaging, and offers important preliminary support for this novel polygenic risk scoring approach. Collectively, these results additionally suggest the possibility that some of the postmortem gene expression changes observed in individuals with MDD may be genetically driven and therefore possibly predate disorder development. Further, these genetically driven changes may bias neural circuit function and associated anhedonia symptoms in similar ways as variants associated with MDD diagnosis in vivo. The fact that T-PRS and PGC-PRS showed both common and distinct patterns of association with brain activity further suggests the two polygenic risk scoring approaches may be complementary and could provide synergistic insight into disorder mechanisms.

Our multivariate PLS analyses associated both T-PRS and the best-fit PGC-PRS with reduced activity to neutral faces across an extensive network of regions, including, but not limited to the corticolimbic circuit (CLC). Specifically, T-PRS was associated with female-specific reduction in activity to neutral faces and male-specific increases in activity to emotional faces and shapes in the mid-frontal and frontal superior regions, supplementary motor area, hippocampus, thalamus, caudate and cerebellum. The PGC-PRS was similarly associated with lower activity to neutral faces in females. While the networks associated with each of the PRS partially overlapped, the PGC-PRS-associated network did not include such a strong response in the frontal and reward regions present in the T-PRS associated network but instead included the insula, postcentral and occipital regions.

Intriguingly, in males with higher T-PRS activity in these same clusters was also blunted in the neutral face condition, although this effect did not reach significance. In contrast, higher T-PRS in males was associated with relatively elevated activity in the shapes and emotional faces conditions. These findings suggest sex-specifictiy of the T-PRS effects, which merits further investigation in future work. Of note, however, only the brain scores associated with the neutral condition were found to be potentially clinically relevant via their association with self-reported anhedonia in women.

For continuity with prior work, we supplemented our multivariate analyses with traditional contrast-based measures focused on the amygdala. Consistent with our multivariate analyses, we found that T-PRS was associated with reduced amygdala reactivity specifically to neutral faces independent of participant sex. Moreover, we demonstrated that the reduced amygdala reactivity to neutral faces mediated the relationship between T-PRS and anhedonia. While the majority of prior work on CLC function in depression has focused on threat-related amygdala reactivity, our results are broadly consistent with recent studies which have found hypoactivity to neutral, but not emotional faces, in youth with severe mood impairment^48^, as well as trait-like reductions in amygdala reactivity to both negative and neutral faces, relative to non-face stimuli, in young adults with seasonal depression^7^. Similarly, reduced CLC reactivity to faces, often accompanied by slower psychomotor speed^24^, is also commonly observed in healthy older adults^41^ or young adults at genetic risk for accelerated brain aging^10, 49^. This phenotypic convergence is consistent with the overlap in molecular pathways implicated in depression and age-related processes, which is well-documented in postmortem gene expression studies^21^ and captured by our T-PRS^11^. Our T-PRS neural reactivity results may thus reflect a genetically driven risk pathway shared between specific depression subtypes, including but not limited to seasonal depression, and both pathological and accelerated aging. No similar relationship was found by the univariate analyses between the PGC-PRS score and amygdala reactivity. However, these univariate ROI-based findings should be interpreted with caution since the T-PRS-based PLS 13 results did not identify the amygdala as a key region and Elliott et al^50^ showed that task-fMRI measures of regional activation have poor reliability in general and that ROIs targeted by the task paradigm are rarely more reliable than non-target ROIs.

Despite the markedly distinct provenance of the PGC-PRS and T-PRS, we found that the two had convergent effects in our multivariate analyses. Both scores were associated with reduced neural reactivity to neutral faces in women across a broader network of regions. This convergence is especially intriguing in light of the fact that the two scores likely reflect distinct biological processes. Specifically, the T-PRS is mainly defined by genes associated with cell signaling and age-related processes such as trophic factor function and cell death^11^, while the independently derived and uncorrelated PGC-PRS is comprised of variants near genes associated with largely non-overlapping biological pathways pertaining to HPA axis hyperactivation, neuroinflammation, and obesity^25^. Collectively, these results suggest that in women, multiple distinct pathways of genetic risk may converge onto a common neural phenotype characterized by blunted neural response to neutral faces. The results are also consistent with the possibility that this neural phenotype may be sensitive to overall genetic risk for depression, independently of any specific underlying molecular mechanism.

Both univariate and multivariate analyses further linked this T-PRS-associated blunting of reactivity to neutral faces with anhedonia, the only depression-specific facet of the MASQ. Furthermore, this association was independent of the remaining three MASQ domains, which capture symptoms unique to anxiety (anxious arousal) or shared between depression and anxiety (general distress scales), and all of these results were independent of early or recent life stress. While prior studies have associated depression with elevated threat-related amygdala reactivity^3–6^, which may partially reflect enhanced neural sensitivity to environmental stress^4^ and may be part of a pathophysiological mechanism shared with comorbid anxiety disorders^51^, the putative novel risk pathway we identify here likely reflects reduced behavioral engagement, which may be depression-specific and relatively independent of experiential factors. These results should be interpreted with caution as the analyses linking brain patterns to mood symptoms were exploratory and therefore uncorrected for multiple comparisons. 14

Nonetheless, the specific link between T-PRS, blunted reactivity to neutral faces and elevated anhedonia emerged in both the univariate and multivariate analyses, giving additional confidence in the potential involvement of T-PRS in an anhedonic pathway of depression risk. Additional caution is warranted in light of the fact that our sample consisted of relatively healthy and highly functioning young adults who likely did not represent the full range of symptom severity. Future studies should aim to test the effects of T-PRS and similar transcriptome-based scores on brain activity in larger clinical populations.

Our results should be interpreted in the context of several limitations. First and foremost, while our PLS analyses accounted for 38–39% of variance shared between PRS and brain activity, effect sizes in our amygdala-focused univariate analyses were relatively small, on the order of 1–2% variance explained. Although these effect sizes are consistent with those seen in other PRS studies using similar or smaller sample sizes^52, 53^, our results suggest multivariate methods leveraging whole-brain voxel-wise co-activation patterns may be better powered to capture complex polygenic effects than analyses focusing on a single region or contrast of interest. Relatedly, task-elicited functional activity in *a priori* regions of interest, including the amygdala, have generally poor reliability^50^ thereby limiting the utility of such measures in individual differences research such as that reported herein^54–57^. Our multivariate analysis may further mitigate this limitation, especially since it models responses to faces and shapes independently of each other, and these independently assessed activity patterns (i.e., shapes or faces vs implicit baseline) have been shown to have higher reliability than difference scores directly contrasting the faces and shapes conditions^58^. Another limitation of this work is that our results are restricted to Non-Hispanic Caucasians. We focused our analysis on this group in order to match the demographic characteristics of the GTEx reference dataset as well as the Ding et al. postmortem dataset, both of which included primarily individuals of European ancestry (85% in GTEx; 97% in Ding et al.^11^). While this decision likely increased our power to detect significant effects, it limits the generalizability of our results to other ethnic groups. Future research using larger multi-ethnic samples for both postmortem and neuroimaging analyses is required to elucidate the functional impact of depression-related transcriptomic changes on brain function across ethnicities.

These limitations notwithstanding, our results introduce a novel tool for bridging depression-associated postmortem molecular gene expression patterns with in vivo brain function and identify a pathway of depression risk associated with reduced neural activity to social stimuli, which may be partially shared with age-related processes. Pending further refinement, this analytic framework bridging postmortem gene expression work with neuroimaging data may offer novel insight into molecular mechanisms underlying MDD or risk thereof and serve to inform future individualized treatment or prevention efforts.

## ACKNOWLEDGEMENTS

YSN is supported by a NARSAD Young Investigator Award from the Brain & Behavior Research Foundation, a Koerner New Scientist Award, and a Paul Garfinkel Catalyst Award administered by the CAMH Foundation. ARH received support from NIH grants R01DA033369 and R01AG049789. ES is supported by NIH Grant R01MH077159 and a NARSAD Distinguished Scientist Award. CH is supported by a NARSAD Young Investigator Award. ANV is currently supported by the Canadian Institutes of Health Research, Canada Foundation for Innovation, the Natioanl Institute of Mental Health, and the CAMH and University of Toronto Foundations.

