## Supplementary figures and images for "Novel polygenic risk score as a translational tool linking depression-related changes in the corticolimbic transcriptome with neural face processing and anhedonic symptoms"

### Supplementary Figure 1

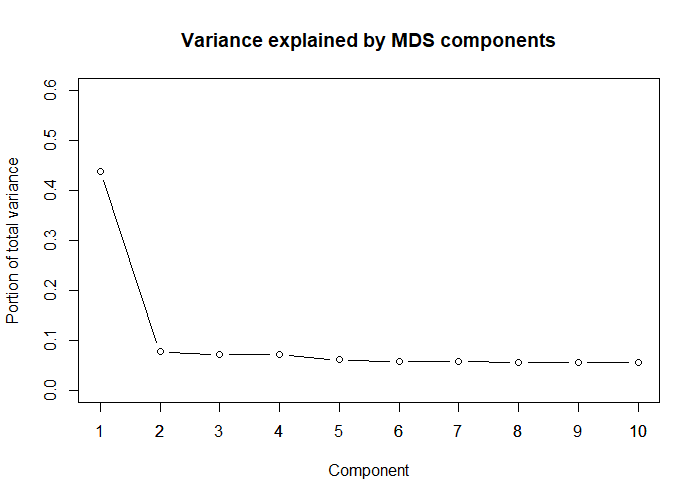

### Supplementary Figure 2

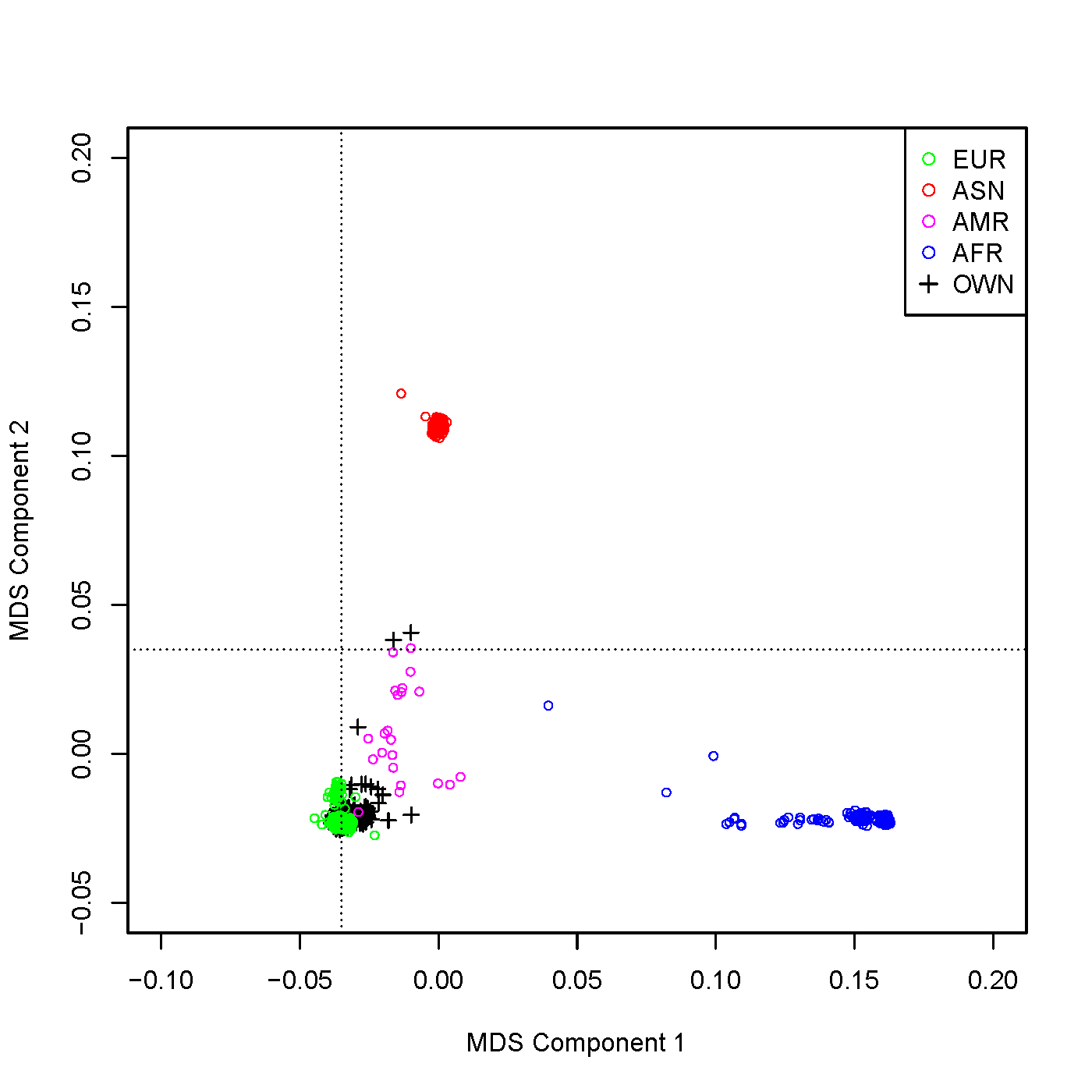

### Supplementary Figure 3

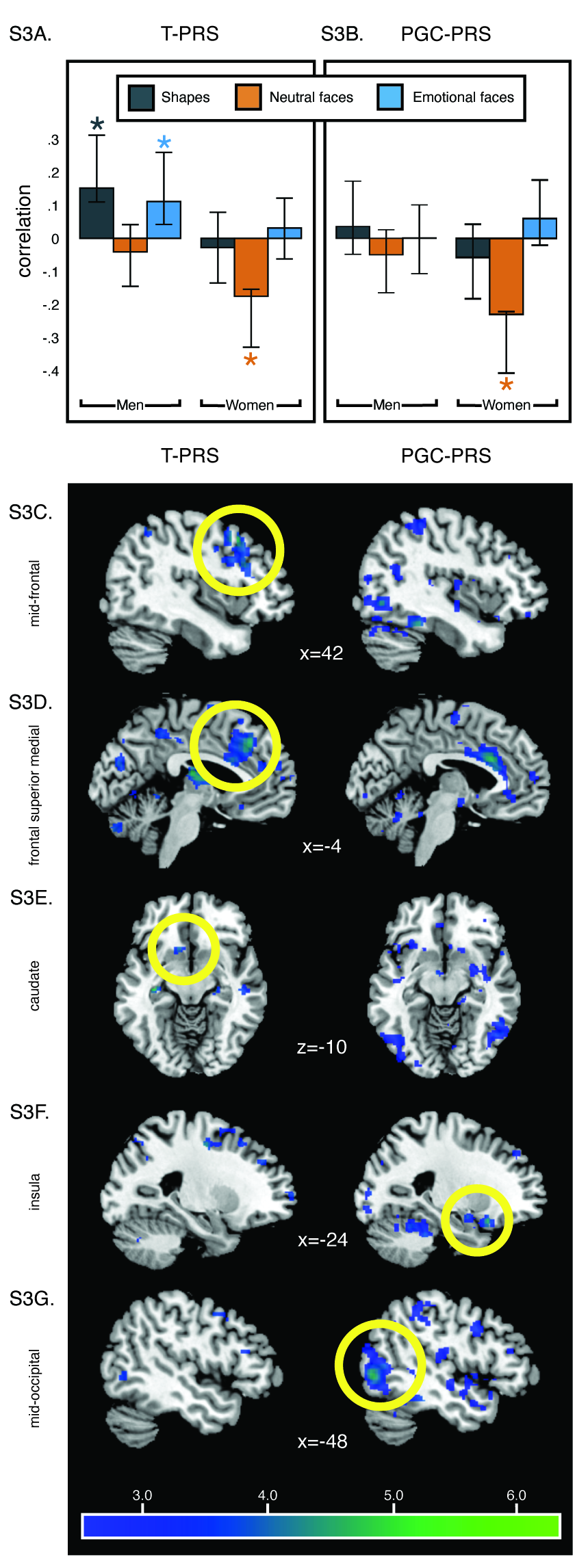

### Supplementary Figure 4

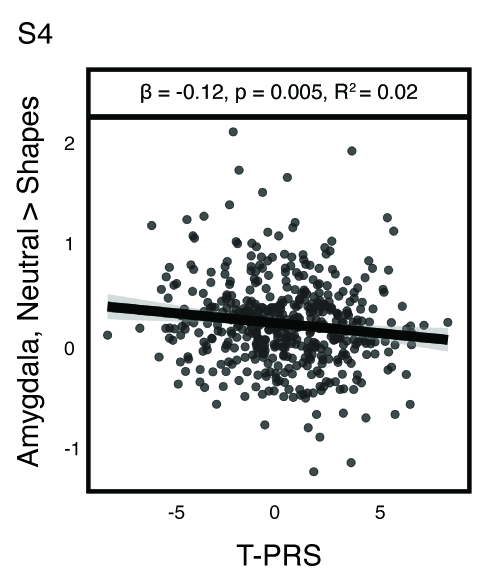

### Supplementary Figure 5

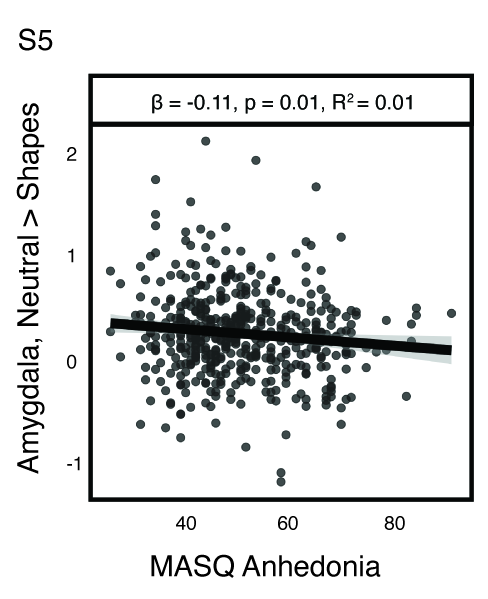

### Supplementary Figure 6

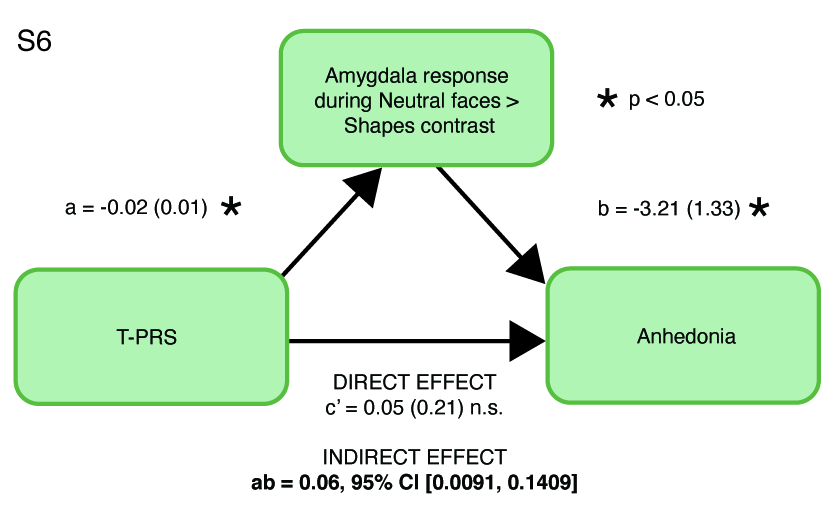
