## Supplementary Methods for "Novel polygenic risk score as a translational tool linking depression-related changes in the corticolimbic transcriptome with neural face processing and anhedonic symptoms"

**SUPPLEMENTARY INFORMATION**

**SUPPLEMENTARY METHODS**

*Acquisition of fMRI data for the amygdala reactivity paradigm*

Functional magnetic resonance imaging (fMRI) during a face matching task was conducted at the Duke-UNC Brain Imaging and Analysis Center using General Electric MR750 3T scanner. The scanner is equipped with high-power, high-duty cycle 50 mT/m gradients at 200 T/m/s slew rate, and an eight-channel head coil for parallel imaging at high bandwidth up to 1 MHz. Global field homogeneity was ensured using a semi-automated high-order shimming program. A series of 34 interleaved axial functional slices aligned with the anterior–posterior commissure (AC–PC) plane were acquired for full-brain coverage using an inverse-spiral pulse sequence to reduce susceptibility artifacts (TR/TE/flip angle = 2000 ms/30 ms/60; FOV = 240 mm; 3.75 × 3.75 × 4 mm voxels; inter-slice skip = 0). Four initial RF excitations were performed (and discarded) to achieve steady-state equilibrium. To allow for spatial registration of each participant's data to a standard coordinate system, high-resolution three-dimensional structural images were acquired in 34 axial slices coplanar with the functional scans (TR/TE/flip angle = 7.7 s/3.0 ms/12; voxel size = 0.9 × 0.9 × 4 mm; FOV = 240 mm, interslice skip = 0).

*fMRI data pre-processing of the amygdala reactivity paradigm*

Preprocessing was conducted using SPM8 ([www.fil.ion.ucl.ac.uk/spm](http://www.fil.ion.ucl.ac.uk/spm)). As described in Carre et al^1^, images for each participant were realigned to the first volume in the time series to correct for head motion, spatially normalized into a standard stereotactic space (Montreal Neurological Institute template) using a 12-parameter affine model (final resolution of functional images = 2 mm isotropic voxels), and smoothed to minimize noise and residual differences in gyral anatomy with a Gaussian filter set at 6 mm full width at half maximum. Next, voxel-wise signal intensities were normalized to the whole-brain global mean. Variability in single-subject whole-brain functional volumes was determined using the Artifact Recognition Toolbox (<http://www.nitrc.org/projects/artifact_detect>). Individual whole-brain BOLD fMRI volumes meeting at least one of two criteria were assigned a lower weight in the determination of task-specific effects: (1) significant mean volume signal intensity variation (i.e., within-volume mean signal greater or less than 4 SD of mean signal of all volumes in time series) and (2) individual volumes where scan-to-scan movement exceeded 2 mm translation or 2° rotation in any direction.

*Whole-brain analysis using Partial Least Square regression*

‘Behavioral’ PLS^2^ was run relating patterns of neural activity to polygenic risk scores (T-PRS and PGC-PRS). PLS is a model-free, multivariate analysis approach for relating patterns of brain activity to continuous variables (typically considered a ‘behavioral score’, but in this case, a PRS). Three fMRI task conditions (Neutral Faces, Emotional Faces or Shapes) were entered, along with either a T-PRS or PGC-PRS (analyzed in separate PLS models). Block design PLS was utilized, which calculates cross-covariance between signal amplitude changes during a block of scanning to a behavioral score. The resulting cross-covariance matrix was then decomposed using singular value decomposition to create a set of orthogonal latent variables (LVs) that optimally represent relationships between voxels activity and behavioral scores across each experimental condition (Neutral Faces, Emotional Faces or Shapes). For each LV, a pattern of voxels (the ‘brain pattern’) demonstrates the relationship with activity in the voxel and the ‘design pattern’ represents the correlation between voxel activity and the gene score for each condition. Voxel weight is expressed as salience, which is proportional to the covariance of activity in that voxel and the design pattern expressed by that LV.

Similarly to Hawco et al^3^, we conducted statistical evaluation of each LV using permutation testing. A total of 500 permutations were run. The permutation score is presented as a p-value and determines if the effect represented by the LV is sufficiently strong to be differentiated from random noise. As the permutations are performed at the level of the entire PLS analysis (rather than on individual LVs), multiple comparisons for the number of LVs is not necessary. LV’s were considered significant if the permutation score was less than 0.05. A bootstrapping procedure with 1000 iterations was used to test if specific voxels were reliably related to the LV. A bootstrap ratio for each voxel was calculated as the voxel salience divided by its bootstrap standard error. Bootstrap ratio is presented in each voxel map to, reflecting the reliability of each voxel as a part of the spatial pattern expressed for that LV. A bootstrap ratio of 2.5 (corresponding to > 95% reliability) was used to threshold all voxel pattern maps in PLS.

*Posthoc univariate analyses focused on the amygdala*

After preprocessing, a first-level, single-subject general linear model (GLM) was used to estimate BOLD responses during (1) Emotional/Threat (Angry and Fearful) vs. Shapes, (2) Emotional/Threat vs. Neutral faces, (3) Neutral faces vs. Shapes contrasts. We focused on these three contrasts, because taken together they allow for dissociating aspects of amygdala reactivity specific to threat from those associated with general social novelty^4^. These individual contrast images (i.e., weighted sum of the beta images) were then used in second-level random-effects models to determine mean condition-specific neural reactivity using one-sample t-tests with a voxel-level statistical threshold of p < .05, family-wise error (FWE) corrected for multiple comparisons across the anatomically defined (Automated Anatomical Labeling [AAL] Atlas-based) amygdala. Hemisphere-specific estimates for the three contrasts of interest were extracted from functional clusters exhibiting a main effect of task using Marsbar (version 0.41) and exported to JMP statistical software version 10.0.0 (SAS Institute Inc., Cary, NC) to perform statistical analyses.

First, a single full-factorial repeated-measures GLM assessed the relationships between either PRS (modeled as a continuous between-subject factor), and sex (x2), hemisphere (x2), and contrast (x3) (modeled as within-subject factors) on amygdala reactivity. Post hoc linear regressions then followed any potential interactions. Next, multiple regression assessed the relationships between self-reported mood and anxiety symptoms and amygdala reactivity. Finally, a moderated mediation used a bootstrapping approach to assess the possible mediation of the relationship between T-PRS and depressive symptoms by amygdala reactivity. All GLM were conducted with and without age, C1, early and recent life stress, and MDD diagnosis as covariates.

**SUPPLEMENTARY RESULTS**

*Polygenic risk and amygdala reactivity*

The GLM focusing on amygdala reactivity yielded a two-way interaction between T-PRS and task contrast (F_(2,947)_=6.74, p=0.0012), and no interactions with sex (p>0.14) or hemisphere (p>0.32). Posthoc analyses revealed that T-PRS predicted BOLD response in amygdala reactivity for the contrast of neutral faces > shapes (beta=-0.12, p=0.005, R^2^=0.02; **Supplementary Figure 4**) and this relationship was independent of all covariates (beta=-0.13, p=0.004; all covariates p>0.21). No relationships were present for any contrast involving threat-related emotional expressions (see **Supplementary Table 6**).

Similar GLM using the PGC-PRS score instead of the T-PRS score did not show any significant effects of the PGC-PRS or its interactions with task contrast, sex, or hemisphere on amygdala BOLD response (p>0.06).

*Amygdala reactivity and depressive symptoms*

Consistent with multivariate results, blunted amygdala reactivity to neutral faces was uniquely associated with the anhedonia subscale (beta=-0.11, p=0.01, R^2^=0.01; **Supplementary Figure 5**), even when controlling for other MASQ domains (beta=-0.14, p=0.02, R^2^=0.01). As suggested by the above pairwise associations, there was evidence of statistical mediation wherein T-PRS mapped onto anhedonia indirectly through amygdala reactivity to neutral faces (ab = 0.06, SE = 0.03, 95% CI [0.009, 0.141]; **Supplementary Figure 6**).

**SUPPLEMENTARY FIGURES**

**Supplementary Figure 1. MDS scatter plot of the two main components based on filtered genome-wide genotype data.** Each dot represents one individual colored by 1000 Genomes ancestry. Color code: red: Asians; blue: Africans; green: Europeans; pink: Admixed Americans; the black cross represents DNS sample (OWN).

**Supplementary Figure 2. Portion of variance explained by MDS components.**

**Supplementary Figure 3.** **Brain response to social stimuli associated with the novel T-PRS and the PGC-PRS and corrected for C1.** Higher levels of T-PRS were associated with lower activity in the LV1 clusters during the Neutral faces condition in women and higher activity in the LV1 clusters during the Emotional faces and Shapes conditions in men (**2A**). Higher levels of PGC-PRS were associated with lower activity in the LV1 clusters during the Neutral faces condition in women (**2B**).

Specifically, higher levels of T-PRS were more strongly associated with lower activity in frontal regions, including the right mid-frontal (**2C**) and frontal superior medial (**2D**), and caudate (**2E**) and higher levels of PGC-PRS were associated with low activity in insula (**2F**) and mid-occipital (**2G**). These clusters survived the 2.5 bootstrap ratio (corresponding to 95% reliability) and were greater than 20 voxels.

**Supplementary Figure 4.** **Higher T-PRS score predicted more blunted amygdala response during the neutral faces > shapes contrast** (beta=-0.12, p=0.005, R^2^=0.02).

**Supplementary Figure 5**. **Lower response in amygdala to neutral faces was associated with greater self-reported anhedonia** (beta=-0.11, p=0.01, R^2^=0.01).

**Supplementary Figure 6. The relationship between T-PRS and anhedonia was mediated by amygdala reactivity to neutral faces** (ab = 0.06, SE = 0.03, 95% CI [0.009, 0.141]).

**SUPPLEMENTARY TABLES**

**Supplementary Table 1** -**Number of participants in our sample (n=482) meeting criteria for at least one DSM IV Axis I diagnosis.**

| **DSM IV Axis I Diagnosis** | **n** |
| --- | --- |
| Agoraphobia (with/ without history of Panic Disorder) | 12 |
| Alcohol Abuse | 35 |
| Alcohol Dependence | 30 |
| Bipolar disorder (past) | 16 |
| Generalized Anxiety Disorder | 5 |
| Major Depressive Disorder (current or past) | 26 |
| Obsessive Compulsive Disorder | 6 |
| Social Anxiety Disorder | 5 |
| Substance abuse (cannabis) | 13 |
| Substance dependence (cannabis) | 7 |
| **Total** | **114** |

**Supplementary Table 2. List of the 76 genes used for calculation of the transcriptome-based polygenic risk score (T-PRS).**

PRSS3

SYT7

AGA

SPATA7

DTNBP1

MAPK9

SPHK2

SNX24

DGCR2

PPP2R3A

EIF4G3

LXN

KIAA1467

CHERP

ANKRD10

DDT

ARSA

DIDO1

ACOT8

TAF1C

ARMC1

SLC1A1

CCNY

GOSR1

CLCN3

FAM149A

COQ5

NPHP3

KLHL24

KCNIP3

GGCX

ALMS1

IVNS1ABP

PPP3CC

PILRB

DOK4

SIN3B

ATF4

PRKRIP1

ATPIF1

TRAF3

RPA1

FIGNL1

MMACHC

GCC2

BPHL

NEK1

ATIC

TBCD

CIAO1

GALNT13

PIK3R1

RABGEF1

RWDD2B

ALDH4A1

AGL

CAMK2N2

GPR98

RRM1

ZNF558

PPIC

PPM1D

DPY19L1

BRMS1

CSRP2

CPNE7

TMEM86B

TTC3

GAS2L1

RPS26

ADH5

SFI1

WWP2

SFT2D1

HN1L

STARD10

**Supplementary Table 3. Total number of variants included in each of the 9 PGC-PRS scores and the associated cross-block covariance. The scores computed at p_GWAS_<0.001 and p_GWAS_<0.01 passed correction for multiple testing (highlighted in bold). The score at p_GWAS_<0.001 (underlined) fit the data better as reflected in its higher cross-block covariance and was selected for visualization and interpretation.**

| **PGC-PRS** |  | **LV1** |  |  |
| --- | --- | --- | --- | --- |
| **GWAS Threshold** | **Number of SNPs identified** | **Cross-block covariance** |  | **p-value** |
| 0.00000005 | 14 | 0.3821 |  | 0.014 |
| 0.000001 | 33 | 0.3671 |  | 0.022 |
| 0.0001 | 373 | 0.2548 |  | 0.084 |
| **0.001** | **1691** | **0.3792** |  | **0.004** |
| **0.01** | **8744** | **0.3602** |  | **0.004** |
| 0.05 | 27618 | 0.3132 |  | 0.048 |
| 0.1 | 44662 | 0.3645 |  | 0.008 |
| 0.5 | 123069 | 0.3611 |  | 0.018 |
| 1 | 165176 | 0.3591 |  | 0.024 |

**Supplementary Table 4**. **Clusters (n=29) negatively correlated with the T-PRS that survived the 2.5 bootstrap ratio threshold (corresponding to 95% reliability).** Each cluster is mapped to a brain region based on the MNI coordinates of its peak voxel. The anatomical description of the region containing the peak MNI coordinate is based on the Talairach atlas labels. Clusters are ordered based on bootstrap ratio, with the most reliable regions on top of the table.

| **Peak region** | **Hemisphere** | **Peak MNI coordinate** | **Peak bootstrap ratio** | **Number of voxels** |
| --- | --- | --- | --- | --- |
| Thalamus | R | 6 -16 12 | -6.3411 | 758 |
| Hippocampus | L | -34 -22 -10 | -4.9614 | 26 |
| Cerebelum_Crus2 | L | -12 -86 -32 | -4.8373 | 234 |
| Frontal mid | R | 42 16 46 | -4.7537 | 1441 |
| Frontal sup medial | L | -4 30 40 | -4.674 | 1091 |
| Middle frontal | L | -26 -12 48 | -4.4401 | 246 |
| Cingulum mid | R | 2 -38 44 | -4.2885 | 229 |
| Supp motor area | L | -6 -2 72 | -4.0308 | 56 |
| Caudate | L | -10 16 -10 | -4.0166 | 146 |
| Frontal mid | L | -26 66 4 | -4.0062 | 97 |
| Cerebelum_Crus1 | R | 12 -82 -28 | -3.9982 | 146 |
| Parietal inf | R | 44 -54 54 | -3.9619 | 129 |
| Precuneus | L | -12 -40 6 | -3.8912 | 22 |
| Parietal inf | L | -30 -48 46 | -3.8539 | 30 |
| Frontal mid | L | -40 14 36 | -3.8294 | 161 |
| Cuneus | L | -2 -80 22 | -3.8255 | 112 |
| Frontal sup | R | 22 66 6 | -3.77 | 20 |
| Frontal sup medial | L | 2 46 40 | -3.7231 | 33 |
| Frontal sup medial | L | 0 62 22 | -3.6669 | 47 |
| Frontal inf tri | L | -50 26 18 | -3.6633 | 21 |
| Precuneus | L | -10 -62 50 | -3.6535 | 34 |
| Cerebelum_Crus2 | L | -36 -74 -36 | -3.6436 | 140 |
| Frontal mid | R | 36 52 8 | -3.6333 | 40 |
| Frontal sup | L | -22 40 34 | -3.599 | 58 |
| Frontal sup medial | L | -2 42 22 | -3.589 | 78 |
| Mid Temporal | R | 48 -22 -10 | -3.5363 | 30 |
| Frontal mid | R | 24 32 40 | -3.4892 | 47 |
| Frontal sup medial | L | 2 64 12 | -3.4288 | 31 |
| Parietal sup | L | -26 -74 52 | -3.3494 | 23 |

**Supplementary Table 5**. **Clusters (n=94) negatively correlated with the PGC-PRS that survived the 2.5 bootstrap ratio (corresponding to 95% reliability).** Each cluster is mapped to a brain region based on the MNI coordinates of its peak voxel. The anatomical description of the region containing the peak MNI coordinate is based on the Talairach atlas labels. Clusters are ordered based on bootstrap ratio, with the most reliable regions on top of the table.

| **Peak region** | **Hemisphere** | **Peak MNI coordinate** | **Peak bootstrap ratio** | **Number of voxels** |
| --- | --- | --- | --- | --- |
| Cingulum_Ant | L | 0 18 29 | -4.9297 | 11225 |
| Amygdala | R | 21 -4 -16 | -4.8801 | 5652 |
| Temporal Inf. | R | 45 -46 -20 | -4.7602 | 14116 |
| Occipital_Mid | L | -48 -72 0 | -4.6198 | 10906 |
| Postcentral | R | 65 -16 17 | -4.5589 | 2328 |
| Insula | L | -24 16 -19 | -4.5311 | 5987 |
| Cerebelum 6 | L | -35 -50 -24 | -3.9891 | 6320 |
| Temporal_Mid | L | -56 -46 4 | -3.9432 | 4102 |
| Temporal_Mid | R | 49 -42 0 | -3.9158 | 750 |
| Cerebelum - Vermis 8 | R | 4 -60 -27 | -3.6913 | 674 |
| Temporal_Pole_Sup | R | 56 12 -8 | -3.6199 | 997 |
| Postcentral | R | 29 -44 65 | -3.6093 | 168 |
| Sub-lobar extra-nuclear | L | -7 -6 -11 | -3.5902 | 180 |
| Frontal_Mid | R | 49 23 33 | -3.5795 | 763 |
| Cerebelum - Culmen | R | 9 -40 -28 | -3.5628 | 238 |
| Temporal_Inf | L | -59 -32 -20 | -3.5473 | 130 |
| Frontal_Inf_Tri | R | 57 16 21 | -3.5381 | 524 |
| Medial Frontal Gyrus | R | 20 -2 53 | -3.5258 | 498 |
| SupraMarginal | R | 33 -38 45 | -3.5036 | 2700 |
| Cerebelum 6 | L | -20 -64 -23 | -3.4571 | 540 |
| Supp_Motor_Area | L | -8 -2 72 | -3.453 | 148 |
| Parietal_Inf | L | -47 -40 40 | -3.4123 | 658 |
| Precentral | R | 48 6 29 | -3.396 | 496 |
| Calcarine | R | 17 -100 -3 | -3.3738 | 186 |
| Temporal_Mid | R | 48 -56 17 | -3.3676 | 1036 |
| Temporal_Sup | R | 53 -18 -4 | -3.3018 | 478 |
| Frontal_Sup_Medial | L | -7 58 12 | -3.2861 | 623 |
| Frontal_Sup_Medial | R | 9 62 25 | -3.2733 | 321 |
| Supp_Motor_Area | R | 9 14 68 | -3.268 | 240 |
| Cerebelum - Vermis 8 | L | 0 -74 -36 | -3.2379 | 182 |
| Frontal_Mid_Orb | R | 37 46 -15 | -3.2132 | 170 |
| Frontal_Sup_Medial | R | 4 63 5 | -3.1809 | 709 |
| Frontal_Sup | L | -24 42 41 | -3.1695 | 237 |
| Frontal_Inf_Orb | R | 44 22 -16 | -3.149 | 34 |
| Temporal_Inf | R | 60 -32 -19 | -3.1295 | 112 |
| Precentral | L | -51 6 36 | -3.1207 | 312 |
| ParaHippocampal | R | 17 -30 -11 | -3.1196 | 182 |
| Frontal_Sup_Medial | R | 8 47 33 | -3.1078 | 88 |
| Frontal_Sup_Orb | R | 20 39 -16 | -3.0763 | 100 |
| Frontal_Sup | L | -19 14 60 | -3.0646 | 260 |
| Occipital_Sup | R | 25 -70 45 | -3.0471 | 146 |
| Postcentral | L | -43 -14 53 | -3.0451 | 182 |
| Postcentral | L | -52 -20 13 | -3.0283 | 220 |
| Angular | R | 61 -52 32 | -3.0082 | 588 |
| Frontal_Sup | R | 21 8 64 | -3.0069 | 260 |
| Calcarine | L | -8 -88 -7 | -2.9822 | 166 |
| Frontal_Inf_Orb | R | 40 31 -4 | -2.9688 | 128 |
| Frontal_Mid | L | -35 22 56 | -2.9621 | 21 |
| Temporal_Sup | R | 60 -34 12 | -2.9616 | 124 |
| Lingual Gyrus | R | 4 -84 -15 | -2.943 | 56 |
| Corpus Callosum | Interhem. | 1 -14 21 | -2.9248 | 192 |
| Frontal_Inf_Oper | L | -43 6 20 | -2.9169 | 78 |
| Frontal_Inf_Tri | L | -40 34 16 | -2.8758 | 188 |
| Cuneus | R | 13 -92 13 | -2.8695 | 52 |
| Frontal lobe | L | -19 -12 60 | -2.858 | 36 |
| Rolandic_Oper | R | 48 -30 17 | -2.8218 | 220 |
| Temporal_Sup | L | -63 -16 5 | -2.8215 | 146 |
| Precentral | R | 36 -26 61 | -2.8172 | 84 |
| Frontal_Sup_Medial | L | -8 67 4 | -2.8021 | 110 |
| Frontal_Mid | L | -40 27 45 | -2.7993 | 39 |
| Frontal_Mid_Orb | R | 32 55 -3 | -2.7954 | 181 |
| Cerebelum_Crus1 | R | 29 -86 -28 | -2.7792 | 34 |
| Insula | R | 29 16 -16 | -2.7611 | 86 |
| Frontal_Sup | L | -11 43 33 | -2.7567 | 76 |
| Occipital_Sup | R | 24 -68 20 | -2.7529 | 102 |
| Olfactory | R | 5 18 -12 | -2.7488 | 87 |
| Temporal_Mid | R | 65 -38 1 | -2.7418 | 190 |
| Sub-lobar extra-nuclear | L | -20 19 -8 | -2.7173 | 52 |
| Frontal_Inf_Oper | R | -35 4 28 | -2.6993 | 32 |
| Cingulate Gyrus | L | -15 -20 37 | -2.6952 | 56 |
| Caudate | R | 17 -8 25 | -2.6922 | 36 |
| Cerebelum_4_5 | L | -7 -60 -7 | -2.6901 | 124 |
| Frontal_Mid | L | -27 46 32 | -2.681 | 52 |
| Parietal_Sup | R | 16 -66 56 | -2.6785 | 108 |
| Cingulum_Mid | L | -3 -8 40 | -2.6766 | 76 |
| Superior Temporal Gyrus | R | -35 2 -19 | -2.6718 | 36 |
| Temporal_Sup | R | 56 -16 1 | -2.6671 | 134 |
| Lingual | R | 9 -74 -11 | -2.6586 | 70 |
| Parietal_Sup | L | -19 -72 57 | -2.6443 | 42 |
| Cerebelum_Crus1 | R | 9 -86 -24 | -2.6429 | 50 |
| Lingual | L | -20 -52 -4 | -2.6403 | 52 |
| Caudate | L | -8 8 0 | -2.6359 | 40 |
| Cerebelum_Crus1 | L | -36 -76 -20 | -2.6256 | 26 |
| Insula | L | -35 -18 1 | -2.6241 | 48 |
| Lingual | R | 9 -60 -3 | -2.624 | 60 |
| Temporal_Mid | L | -51 4 -23 | -2.6143 | 24 |
| Cerebelum - Culmen | R | 4 -54 -12 | -2.6081 | 38 |
| Postcentral | R | 17 -38 69 | -2.6061 | 30 |
| Temporal_Pole_Sup | L | -36 12 -27 | -2.5969 | 42 |
| Temporal_Sup | L | -64 -36 12 | -2.5925 | 34 |
| Cingulum_Mid | R | 1 -20 33 | -2.5852 | 36 |
| Frontal_Sup | R | 17 35 45 | -2.5778 | 36 |
| Frontal_Inf_Orb | R | 25 19 -19 | -2.5584 | 27 |
| Caudate | L | -4 8 -7 | -2.557 | 42 |

**Supplementary Table 6.** **The effect of T-PRS on BOLD response in amygdala.** Significant effects are shown in bold font.

| **Contrast** |  |
| --- | --- |
|  | **PRS** |
| Emotional vs. Neutral faces | beta=0.06, p=0.20 |
| Emotional faces vs. Shapes | beta=0.06, p=0.20 |
| **Neutral faces vs. Shapes** | **beta=-0.13, p=0.005** |
